# Habitat complexity and benthic predator-prey interactions in Chesapeake Bay

**DOI:** 10.1101/224089

**Authors:** Cassandra N. Glaspie, Rochelle D. Seitz

## Abstract

In Chesapeake Bay, the soft-shell clam *Mya arenaria* (thin-shelled, deep-burrowing) exhibits population declines when predators are active and persists at low densities. In contrast, the hard clam *Mercenaria mercenaria* (thick-shelled, shallow-burrowing) has a stable population and age distribution. We examined the potential for habitat and predators to control densities and distributions of bivalves in a field caging experiment (*Mya* only) and laboratory mesocosm experiments (both species). In the field, clams exposed to predators experienced 76.3% greater mortality as compared to caged individuals, and blue crabs were likely responsible for most of the mortality of juvenile *Mya*. In mesocosm experiments, *Mya* had lower survival in sand and seagrass than in shell hash or oyster shell habitats. However, crabs often missed one or more prey in seagrass, shell, and oyster shell habitats. Predator search times and encounter rates declined when prey were at low densities, likely due to the added cost of inefficient foraging; however, this effect was more pronounced for *Mya* than for *Mercenaria*. *Mercenaria* had higher survival than *Mya* in mesocosm experiments, likely because predators feeding on *Mercenaria* spent less time foraging than those feeding on *Mya*. *Mya* may retain a low-density refuge from predation even with the loss of structurally complex habitats, though a loss of habitat refuge may result in clam densities that are not sustainable. A better understanding of density-dependent predator-prey interactions is necessary to prevent loss of food-web integrity and to conserve marine resources.

## INTRODUCTION

Predators exhibit top-down control on communities, influencing the abundance, size structure, and distribution of prey by restricting their survival or activity in time and space [1–3]. Predators also influence community function by preying upon dominant species [4–6]. To understand the structure and function of a community, it is important to consider the impact of the predators. Prey populations experience the effects of predation differently depending on how abundant the prey species is and, for actively foraging predators, how quickly the predator can find and consume prey [7]. The degree to which a predator can reduce prey abundance is a function of the probability of encountering a prey item, and the probability that the prey item will be eaten, given that it has been encountered. Both factors depend on the characteristics of the prey, the predator, and other environmental factors [6].

Bivalve mollusks exhibit a number of morphological and behavioral characteristics to defend against predators. Armor and aggregation decrease rates of predation, allowing predators and prey to coexist in the same space. For example, the infaunal, shallow-burrowing, hard-shell clam *Mercenaria mercenaria* (hereafter, *Mercenaria*) has a relatively thick shell that protects it from predation by blue crabs *Callinectes sapidus*; clams larger than 40 mm cannot be crushed and therefore coexist with crabs [8]. Other bivalves must avoid predators to survive; the shell of a soft-shell clam *Mya arenaria* (hereafter, *Mya*) is thin and has a permanent gape, indicating that for this species, shell thickness is not an important mode of protecting against attack by predators [9]. To avoid predation, large individuals of *M. arenaria* achieve a non-coexistence refuge by burrowing 25-30 cm deep in the sediment, out of range of foraging predators, which rarely consume clams buried deeper than 10 cm [10].

Habitat also plays an important role in predator defense strategies of marine bivalves. Predators in habitats that are not complex have a greater effect on prey than those in complex habitats [11,12]. Vegetated or shell habitat provides a refuge from predation for many prey [12,13], and increased sediment grain size allows infaunal species to avoid predators more effectively than in fine sediments [10,14,15]. Complex habitats increase metabolic costs associated with foraging, and as these costs become too high, predators may opt to conserve energy or forage elsewhere [16,17].

The functional response is a way to quantify predator foraging efficiency [7]. A predator’s functional response is the relationship between the number of prey consumed per predator and prey density [18]. Predators that search for prey exhibit a density-dependent functional response, because the encounter rate depends on prey density. In a type II density-dependent response, handling rate and attack rate remain constant as prey density increases [7]. Prey consumed per predator increases with increasing prey density, but the rate of increase declines to an upper asymptote. The asymptote is reached when the predator becomes satiated and spends less time foraging, or when the predator is limited by the amount of time it takes to consume prey [7]. A type III sigmoidal density-dependent response occurs when a predator becomes more active as prey density rises, which means attack rate is a function of prey density [7]. Type II and type III functional responses are very different biologically, since type III functional responses create a refuge for prey at low densities, which may result in prey persistence over time, even if a population is driven to low abundance [7,19,20].

The main parameters in a functional response model are encounter rate and handling time [7], both of which change as a function of prey mortality, prey behavior, and habitat type. For the purposes of this study, the encounter rate was defined as the number of encounters with prey divided by the amount of time a predator spends foraging, or actively looking for prey; and the handling time was defined as the amount of time a predator spends manipulating or eating a prey item. For thick-shelled bivalves, the consumption rate of their predators is determined more by handling time than encounter rate; in this case, a type II functional response is more likely [14]. For burrowing, thin-shelled bivalves, encounter rate is more important than handling time for their predators [2], which means that a density-dependent sigmoidal (type III) response is likely [14]. The biological mechanism behind a type III response is that low encounter rates often lead to low activity levels or predators emigrating from the area [21]. The functional response of a predator-prey interaction can also be habitat specific. Reduced sediment penetrability [14] or increased vegetative cover [22] may lead to decreased encounter rate, and this may change the functional response by creating or strengthening a low-density refuge from predation. The functional response also changes with ontogeny, as small bivalves may not have sufficiently thick shells to impact predator handling time or burrow deeply enough to reduce encounter rate with predators [23].

In the Chesapeake Bay, two commercially valuable clam species, the soft-shell clam *Mya* and the hard clam *Mercenaria* have very different population dynamics. Adult and sub-adults of *Mya* exist in the Bay at low abundance except immediately after spring recruitment, and juveniles are nearly completely consumed by predators each year [24] (Fig 1). *Mercenaria* is fairly abundant throughout the year, and all size classes persist in the Bay in all seasons [25] (Fig 1). The different dynamics of these species may be due to predator-prey dynamics, since the two species exhibit different predator-avoidance strategies. Specifically, the persistence of *Mya* at low abundance may be due to a low-density refuge, especially in complex habitats that prevent efficient foraging by the species’ main predators, the blue crab *Callinectes sapidus* [26,27] and the cownose ray *Rhinoptera bonasus* [28].

**Fig 1.**
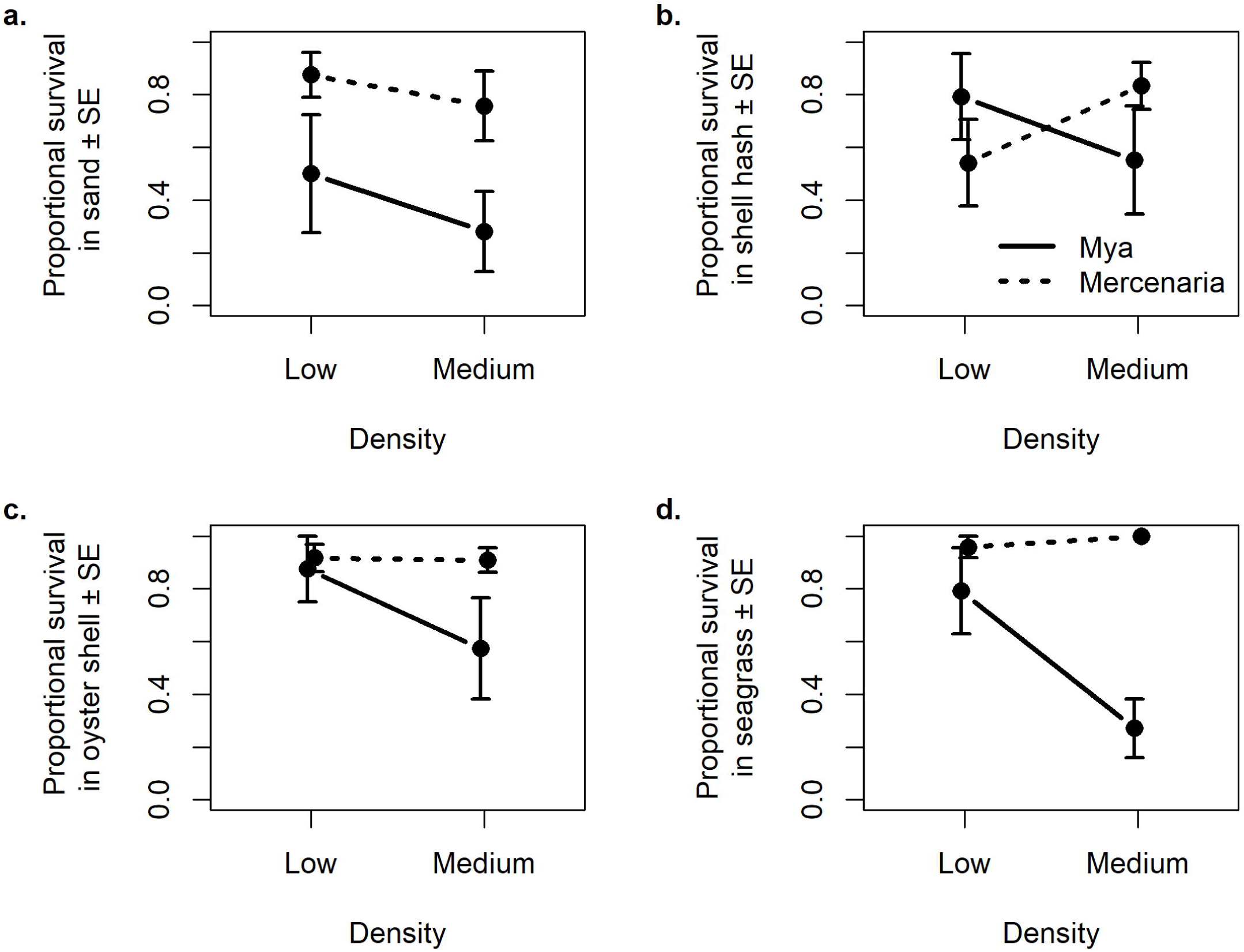
Size frequency histograms of *Mercenaria mercenaria* (left) and *Mya arenaria* (right) in lower Chesapeake Bay. Samples were collected in spring (a-b), summer (c-d), and fall (e-f) for two years starting in fall 2011. Sizes expressed are biomass (g AFDW) for *Mercenaria* [25] and *Mya* [24].

This study aims to examine the nature of blue crab-bivalve predator-prey interactions for these two infaunal bivalves, including the role of structural refuge (in the form of complex habitat) on these interactions, using both field and laboratory experiments. In field caging experiments, we hypothesized the following: 1) blue crabs and cownose rays are both sources of mortality for sub-adult *Mya* (evidenced as a significant difference in *Mya* survival among all caging treatments); and 2) the presence of seagrass increases clam survival rates as compared to sand and mud (for all plots without a complete cage). In laboratory mesocosm experiments, we hypothesized the following: 1) predators on sub-adult *Mya* exhibit a type III functional response and predators on sub-adult *Mercenaria* exhibit a type II functional response (evidenced as a significant species-density interaction); 2) complex (as compared to unstructured) habitats increase the extent of the low-density refuge for species using density as a refuge, which manifests as increased proportional survival in complex habitats as compared to sand, but only for *Mya* (evidenced as a significant species-habitat interaction); 3) *Mercenaria*’s armor leads to increased handling time compared to *Mya* (evidenced as a significant main effect of species on handling time); 4) low densities, complex habitat, and deep-burrowing prey result in decreased blue crab search time, due to the added cost of inefficient foraging (evidenced as a 3-way interaction between species, density, and habitat), and 5) there is a decreased encounter rate at low densities of *Mya* compared to high densities (evidenced as a significant species-density interaction).

## MATERIALS AND METHODS

### Field caging experiment

A caging study was conducted in patchy seagrass, sand, and mud near-shore habitats (1.5-2 m depth mean high water) in May 2014 near the mouth of the York River, VA (between 37.258323, -76.428047 and 37.275197, -76.370150). These habitat types represented decreasing habitat complexity from seagrass to mud; compared to mud, sand provides additional habitat complexity for infaunal bivalves such as *Mya*, altering the functional response [29]. Ten replicate 0.25 m^2^ plots were randomly assigned one of three caging treatments in each habitat: full cage, stockade, or uncaged. Full cages were constructed of 13-mm galvanized wire mesh with PVC frames (0.6 m height, 0.5 m width, 0.5 m length) and were sunk into the sediment approximately 10 cm and secured with PVC legs sunk an additional 30-40 cm. Stockades were constructed by placing 8 10-ft PVC poles around an otherwise unprotected plot at 25-cm intervals. Stockades kept cownose rays out of the plots, while still allowing for crab and fish predation. Uncaged plots were marked with two PVC poles on the diagonals.

Juvenile soft-shell clams (*Mya*) 20-40 mm shell length (mean 28.48 ± 4.41 mm SD) were collected from the York River and held in flow-through tanks until experimentation. Clams were marked individually with permanent marker and transplanted towards the center of the plot at densities of 12 clams per plot (48 m^-2^) [30]. A cage was placed over all transplanted clams to allow them to acclimate overnight and achieve a stable burrowing depth as in previous laboratory experiments under similar temperatures [21], and acclimation cages were removed from stockade and uncaged treatments. After 5 d, the contents of all plots were collected to a depth of 40 cm using a suction sampler [20]. Remaining bivalves were counted and shell fragments were noted as evidence of crab predation. Partial cages were not used to control for caging artifacts due to the short nature of this study and the tendency for partial cages to attract blue crabs. Given the relatively large aperture of the cage mesh (13 mm), we would not expect notable differences in cage artifacts among habitat types over the 5-day trial. Only one density was used in this study due to the presence of wild *Mya* in the area, and the consequent logistical difficulties associated with creating reliable densities.

Proportional survival data were Box-Cox transformed (λ = 0.51) to achieve normality and homogeneous variance (assessed using quantile-quantile and residual plots), and analyzed using two-way ANOVA, with cage type (3 levels: full cage, stockade, and uncaged) and habitat (3 levels: mud, sand, and seagrass) as fixed factors, with α = 0.05 for main effects and α = 0.20 for interaction terms [31]. Post-hoc pairwise comparisons were done using Tukey honest significant difference (HSD) tests. From a pilot caging experiment in 2012, we used a simulation of resampled data to determine that our sample size of n = 10 resulted in the following estimates of statistical power: 1.00 for the main effect of cage type, 0.42 for the main effect of habitat, and 0.87 for the interaction effect.

### Laboratory mesocosm experiment

*Mya* (thin-shelled, deep infaunal) and *Mercenaria* (thick-shelled, shallow infaunal) were exposed to blue crab *C. sapidus* predation in mesocosm tanks of 0.87 m diameter and 0.59 m height, which were partitioned with corrugated plastic to form a rectangular experimental arena (40 cm x 70 cm). Sand was added to the tank to 25 cm depth, and an additional 25 cm of the tank was filled with filtered water from the York River. An aquarium heater held tank temperature constant at 26-27 °C, typical of shallow York River water in the summer months [32], and the water was aerated by air stones placed outside the experimental arena. Trials were randomly assigned one of four habitat treatments: sand alone, sand/shell hash, sand/oyster shell, or sand/seagrass. For trials receiving shell or oyster shell, a constant volume of 0.5-L crushed shell hash (lightly crushed Baltic clam *Macoma balthica*, ribbed mussel *Geukensia demissa*, and *Mercenaria* shell halves) or oyster shell halves was added to the center of the mesocosm tank. Eelgrass (*Zostera marina*) and widgeongrass (*Ruppia maritima*) shoots and rhizomes were collected from the York River and used to construct seagrass mats for use in trials receiving seagrass. Seagrass mats were constructed with 0.5 liter of natural seagrass blades tied onto plastic 1-cm Vexar mesh meant to simulate a rhizome mat. Holes measuring approximately 25 cm^2^ were cut approximately every 10 cm to allow crabs to forage for clams buried under the simulated seagrass mat. The mesh and attached seagrass roots were placed in the center of the tank and completely covered with sand.

Juvenile *Mya* 20-40 mm shell length were collected from the York River and held in flow-through tanks until experimentation. Hard clams *Mercenaria* 30-40 mm shell length were obtained from Cherrystone Aqua-Farms in Virginia. Only hard clams with shell lengths < 40 mm were used in the study, because blue crabs are able to consume clams of this size [33]. Bivalves were placed in the sediment siphon up, away from the edge of the tank to avoid edge effects, and allowed 24 h to achieve a stable burial depth [21]. Each species was transplanted at two densities as determined from the literature, one low and one medium density. When number of prey consumed is converted to proportion of prey eaten per predator, two densities (low and medium) are sufficient to determine whether a low-density refuge exists (positive relationship between proportional mortality and prey density, indicating at type III functional response) or does not exist (negative relationship between proportional mortality and prey density, indicating at type II functional response, as in previous studies [21,34]. Low densities for both species were 4 clams per tank, and medium densities were 11 clams per tank for *Mercenaria* and 16 clams per tank for *Mya* [16,34].

*Callinectes sapidus* were collected from the York River via baited crab pot. All crabs were acclimated to the lab for 1 week or longer and fed fish or clam meat three times per week. At the start of the experiment, one adult male blue crab with a carapace width ≥ 100 mm was added to each tank receiving a predator treatment. Bivalves were exposed to blue crab predation for 48 h, as is common for similar mesocosm studies [20]. Remaining bivalves were excavated and counted upon termination of the experiment. There were six replicates of each habitat/density combination, as well as an equal number of mesocosms set up without predators, which served as controls (though only 0.6% of clams died in predator-free controls and they are not analyzed or discussed further).

Proportional survival data were Box-Cox transformed (λ = -0.14) to achieve normality and homogeneous variance (assessed using quantile-quantile and residual plots), and they were analyzed using three-way ANOVA, with density (2 levels: low and medium), species (2 levels: *Mya* and *Mercenaria*) and habitat (4 levels: sand, shell hash, oyster shell, and seagrass) as fixed factors, with α = 0.05 for main effects and α = 0.20 for interaction terms [31]. Effect size and standard error estimates from a previously conducted mesocosm experiment [21] were used to calculate power to see a significant main effect of density, which was 0.95 for n = 6. Post-hoc pairwise comparisons were done using Tukey HSD tests.

It was not possible to use a different crab for each trial due to space requirements, nor was it possible to use each crab the same number of times due to losses throughout the experiment. Crabs were used between one and five times, and crabs were randomly assigned to trials so there was no bias inherent in the re-use of crabs. An ANCOVA including density, species, habitat, individual crab identity (51 levels), number of times a crab was used (continuous, 1-5), tank (4 levels), and day of the experiment (continuous, standardized using z score transformation) as covariates indicated that there was no difference in proportion of bivalves eaten based on crab identity (F_49, 24_ = 1.23, p = 0.30), number of times the crabs were used (F_1, 24_ = 1.56, p = 0.22), tank (F_3, 24_ = 0.48, p = 0.70), or day of the experiment (F_1, 24_ = 1.15, p = 0.29). These results provided no evidence that crabs exhibited learning behavior, and no evidence for tank effects or trends through time; thus, each trial was treated as an independent replicate.

For half of the trials (n = 3 for each treatment) predator behavior was recorded using an infrared-sensitive camera system. A red spotlight was used to improve night-time video quality without disrupting crab behavior [35]. Videos were used to calculate search time, encounter rate, and handling time. Search time (h) was defined as the total time spent exhibiting foraging behavior, such as probing the sediment with legs or claws or lifting items to mouthparts. Encounter rate (hr^-1^) was defined as the number of encounters (picking up bivalve) divided by the search time. Handling time (h) was defined as the total time spent manipulating or eating a bivalve, divided by the number of encounters. Handling time, search time, and encounter rate were fourth-root transformed to achieve homogeneity and compared for the two bivalve species in different habitat treatments and at different densities using three-way ANOVAs with the same factors as were used for analysis of proportional survival. Post-hoc pairwise comparisons were done using Tukey HSD tests.

All analyses were completed using R statistical software [36], and data and R code files are available in the Knowledge Network for Biocomplexity (KNB) repository [37].

## ETHICS STATEMENT

Virginia Institute of Marine Science is statutorily mandated as Virginia’s scientific advisor on marine- and coastal-related natural resources and exempt from having to obtain a scientific collection permit for non-protected species in Virginia’s waters.

## RESULTS

### Field caging experiment

Over the 5-day caging experiment, mean water temperature at the nearby YKTV2 weather buoy was 18.76 °C (± 1.63 SD). All replicates (n = 10) for the stockade and uncaged plots lasted through the experiment and were subsequently sampled. At least one of the caged plots was lost from each habitat, leaving n = 9 replicates in mud, n = 7 replicates in sand, and n = 8 replicates in seagrass.

As compared to full cages, there was a decrease in proportional survival of 75.6% in stockades and 77.0% in uncaged plots (Fig 2), but the effect of one main effect depended on the conditions of the other (Table 1). Stockade and uncaged treatments had similar survival among habitats (p = 1.0). Mud had significantly lower survival than sand (p = 0.002) or seagrass (p = 0.0002). Seagrass and sand had similar survival (p = 0.86). Due to a significant habitat x cage interaction, main effects need to be interpreted with caution (Table 1). The significant habitat x cage treatment interaction was driven by the full cage treatment, which had different patterns of survival than the other caging treatments (Supp. Table 1). Survival of clams in stockades placed in mud was lower than might be expected with just main effects of habitat and cage type (Supp. Table 1).

**Table 1.**
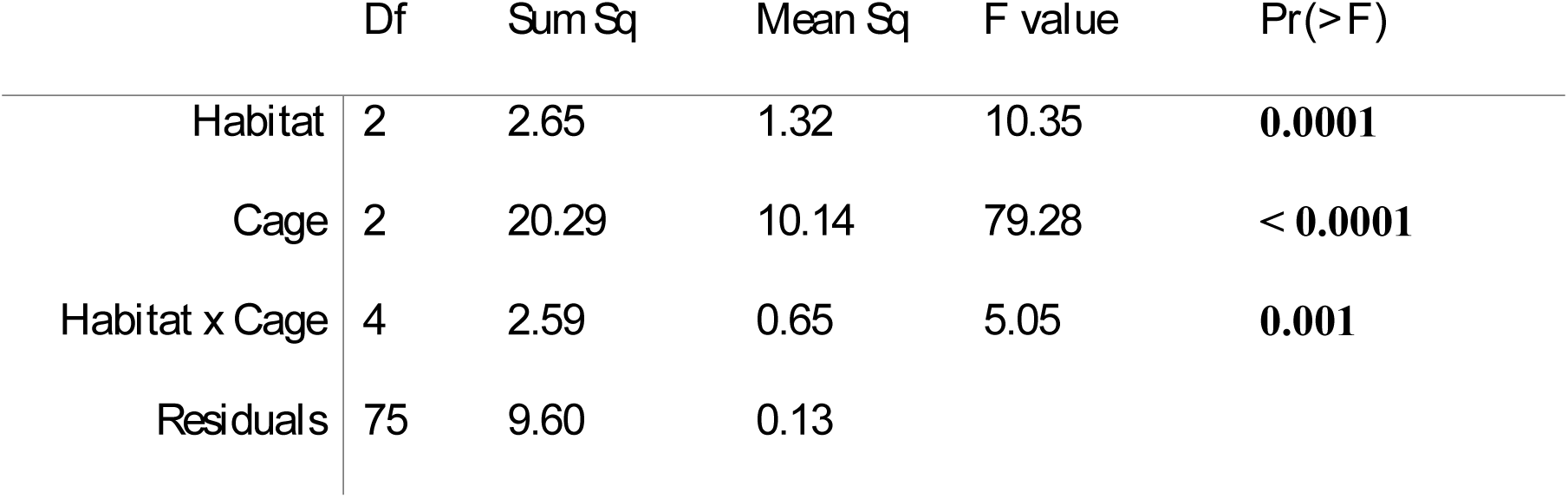
ANOVA summary table for field caging study proportional survival data. Three types of caging treatments (full cage, stockade, and uncaged) were placed in three habitat types (mud, sand, and seagrass); all were included in the ANOVA model as fixed factors. Data were Box-Cox transformed (λ = 0.51) prior to analysis. Significant p values (at α = 0.05 for main effects and α = 0.20 for interaction terms) are bolded.

|  | *Df* | *Sum Sq* | *Mean Sq* | *F value* | *Pr(>F)* |
| --- | --- | --- | --- | --- | --- |
| *Habitat* | 2 | 2.65 | 1.32 | 10.35 | **0.0001** |
| *Cage* | 2 | 20.29 | 10.14 | 79.28 | **< 0.0001** |
| *Habitat x Cage* | 4 | 2.59 | 0.65 | 5.05 | **0.001** |
| *Residuals* | 75 | 9.60 | 0.13 |  |  |

**Fig 2.**
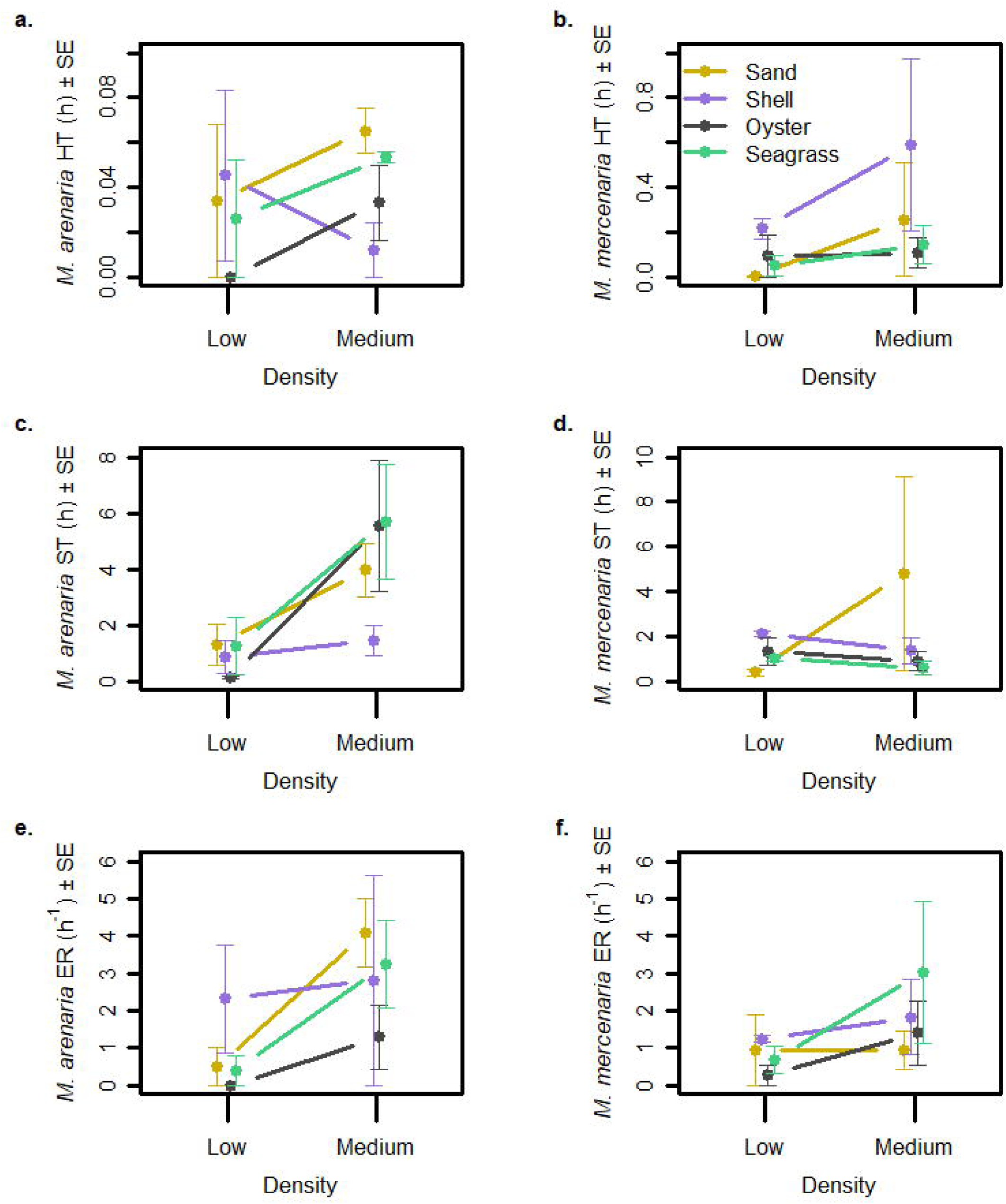
Survival of transplanted juvenile *Mya arenaria* exposed to a natural suite of predators near the mouth of the York River, VA. Shown are mean proportional survival (± 1 SE) after 5 d in the field. Bivalves were placed in full cages (full), stockades, or uncaged plots. Plots were in different habitats (denoted by different color bars). There were n = 10 replicates for the stockade and uncaged plots, and n = 9, 7, and 8 replicates for cages in mud, sand, and seagrass, respectively.

On average, 39.3% of missing clams were recovered as crushed shells within the plots. Mean recovery of crushed shells varied little among caging types and habitats. The highest occurred in stockade plots in sand, with 49.2% (± 28.7 SD) of missing clams recovered as crushed shells, and lowest occurred in uncaged plots in mud, with 24.7% (± 26.5 SD) of missing clams recovered as crushed shells. Not all clams were recovered from caged plots because the suction sampler used to retrieve clams missed some individuals.

### Laboratory mesocosm experiment

In mesocosm experiments, mean proportional survival ranged from 0.27 (*Mya* in seagrass at medium densities) to 1.00 (*Mercenaria* in seagrass at medium densities). Crabs ate at least one *Mercenaria* in 18 out of 48 trials, and ate all offered *Mercenaria* in only one trial (low density in shell). Predation of *Mya* was more common, with at least one *Mya* eaten in 27 out of 48 trials. In the sand at low densities, crabs either ate all of the available *Mya* (occurred 3 times), or none of them (occurred 3 times). In the more-complex habitats (shell hash, oyster shell, and seagrass), crabs offered low densities of clams usually ate none of them (occurred 13 out of 18 trials); only occasionally would a crab eat a portion of the total number of clams offered (1, 2, or 3 clams; occurred 3 times) or all 4 of the clams (occurred 2 times).

*Mya* had significantly lower survival than *Mercenaria* (Fig 3; Table 2), but the effect of one main effect depended on the conditions of the others. There was some evidence that bivalves had lower proportional survival in trials with medium bivalve densities than in trials with low bivalve densities (Table 2). There were no significant differences in survival by habitat type or bivalve density (Table 2), but there were significant species x habitat interactions. *Mya* in medium densities had lower survival than the other species x density combinations, driving a significant species x density interaction (Supp. Table 2). In sand and seagrass, *Mya* had lower survival than some other species x habitat combinations, driving a significant species x habitat interaction (Supp. Table 3).

**Table 2.** ANOVA results for mesocosm study proportional survival of juvenile clams, as well as handling time (HT), search time (ST), and encounter rate (ER) of blue crabs *Callinectes sapidus* feeding on juvenile clams. Two species (*Mya arenaria* and *Mercenaria mercenaria*) were offered to blue crabs *Callinectes sapidus* at two densities (low and medium) in tanks with four different habitats (sand, sand with shell hash, sand with oyster shell halves, and sand with live seagrass); all were included in the ANOVA model as fixed factors. Data were Box-Cox transformed (λ = -0.14; survival only) or fourth-root transformed (HT, ST, and ER) prior to analysis. Significant p values (at α = 0.05 for main effects and α = 0.20 for interaction terms) are bolded.

|  | *Survival* | *HT* | *ST* | *ER* |
| --- | --- | --- | --- | --- |
| *Species* | **F<sub>1,80</sub> = 15.90, p = 0.0001** | F<sub>1,32</sub> = 2.87, p = 0.10 | F<sub>1,32</sub> = 0.69, p = 0.41 | F<sub>1,32</sub> = 0.07, p = 0.79 |
| *Density* | F<sub>1,80</sub> = 3.68, p = 0.06 | **F<sub>1,32</sub> = 4.28, p = 0.05** | **F<sub>1,32</sub> = 10.10, p = 0.003** | **F<sub>1,32</sub> = 6.46, p = 0.02** |
| *Habitat* | F<sub>3,80</sub> = 1.86, p = 0.14 | F<sub>3,32</sub> = 1.23, p = 0.32 | F<sub>3,32</sub> = 0.31, p = 0.82 | F<sub>3,32</sub> = 1.19, p = 0.33 |
| *Species x Density* | **F<sub>1,80</sub> = 7.17, p = 0.01** | F<sub>1,32</sub> = 0.03, p = 0.88 | **F<sub>1,32</sub> = 11.38, p = 0.002** | F<sub>1,32</sub> = 0.95, p = 0.34 |
| *Species x Habitat* | **F<sub>3,80</sub> = 2.19, p = 0.10** | **F<sub>3,32</sub> = 2.01, p = 0.13** | F<sub>3,32</sub> = 1.13, p = 0.35 | F<sub>3,32</sub> = 0.65, p = 0.59 |
| *Density x Habitat* | F<sub>3,80</sub> = 0.65, p = 0.58 | F<sub>3,32</sub> = 0.91, p = 0.45 | F<sub>3,32</sub> = 1.47, p = 0.24 | F<sub>3,32</sub> = 1.27, p = 0.30 |
| *Species x Density x Habitat* | F<sub>3,80</sub> = 0.62, p = 0.61 | F<sub>3,32</sub> = 0.25, p = 0.86 | **F<sub>3,32</sub> = 2.08, p = 0.12** | F<sub>3,32</sub> = 0.54, p = 0.66 |

**Fig 3.**
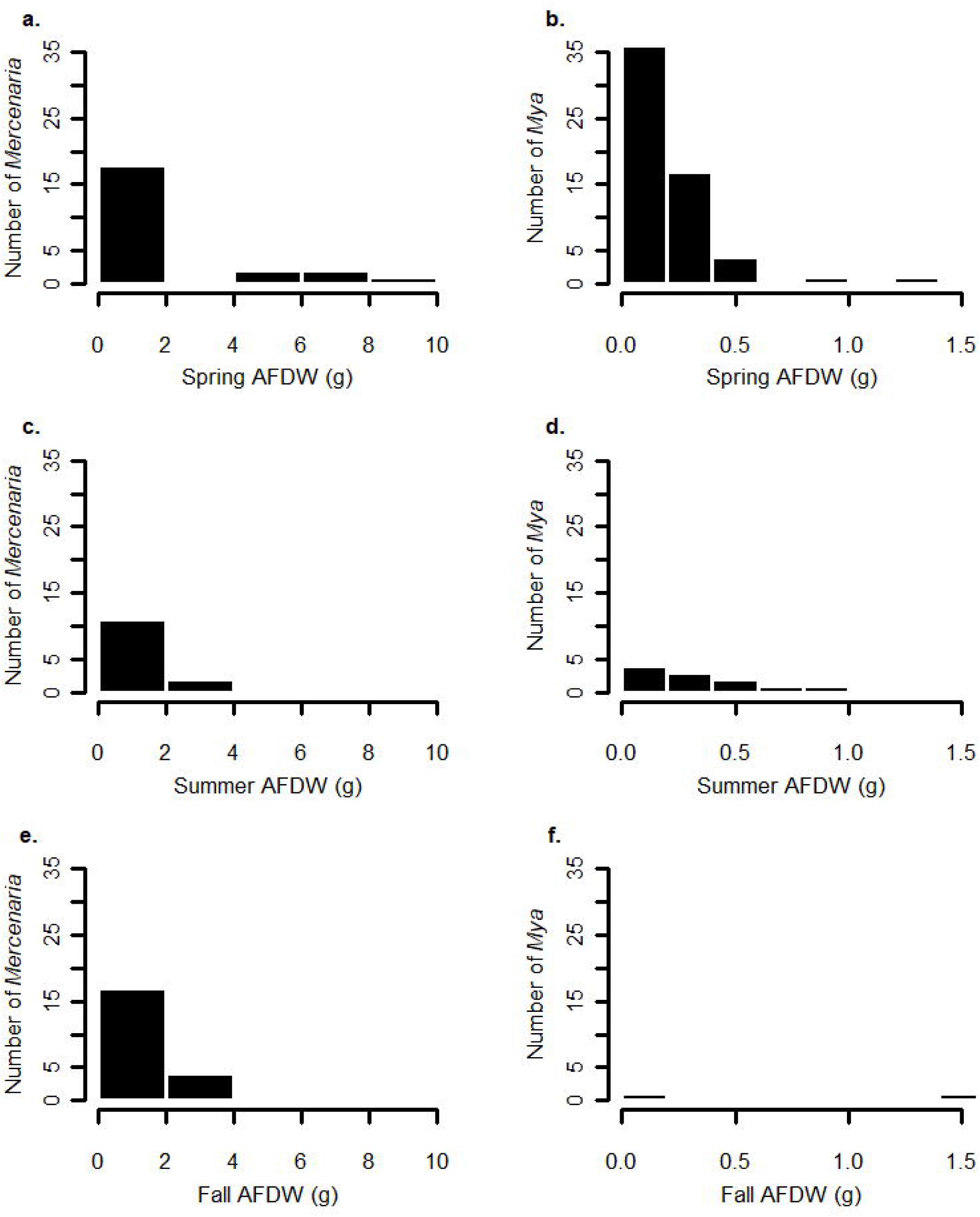
Density-dependent predation in different habitats. Mean juvenile *Mya arenaria* and *Mercenaria mercenaria* proportional survival (± 1 SE) in mesocosms when exposed to blue crab predation in a) sand, b) shell hash, c) oyster shell, and d) seagrass. Solid black lines are mean proportional survival for *Mya* at two initial densities of 4 and 16 per tank, and dashed black lines are mean proportional survival for *Mercenaria* at two initial densities of 4 and 11 per tank.

Handling time was significantly lower in low-density trials than in medium-density trials (Fig 4a, b; Table 2), but the effect of one main effect depended on the conditions of the others. The two treatments with the longest mean handling times were *Mercenaria* at medium density in shell hash (1.31 h) and *Mercenaria* at medium density in sand (0.76 h). All other treatments had mean handling times of 0.30 h or less. The overall mean handling times for *Mercenaria* and *Mya* were 0.18 h and 0.03 h, respectively. In shell hash, *Mercenaria* had longer handling times than the rest of the species x habitat combinations, driving a significant species x habitat interaction (Supp. Table 4).

**Fig 4.**
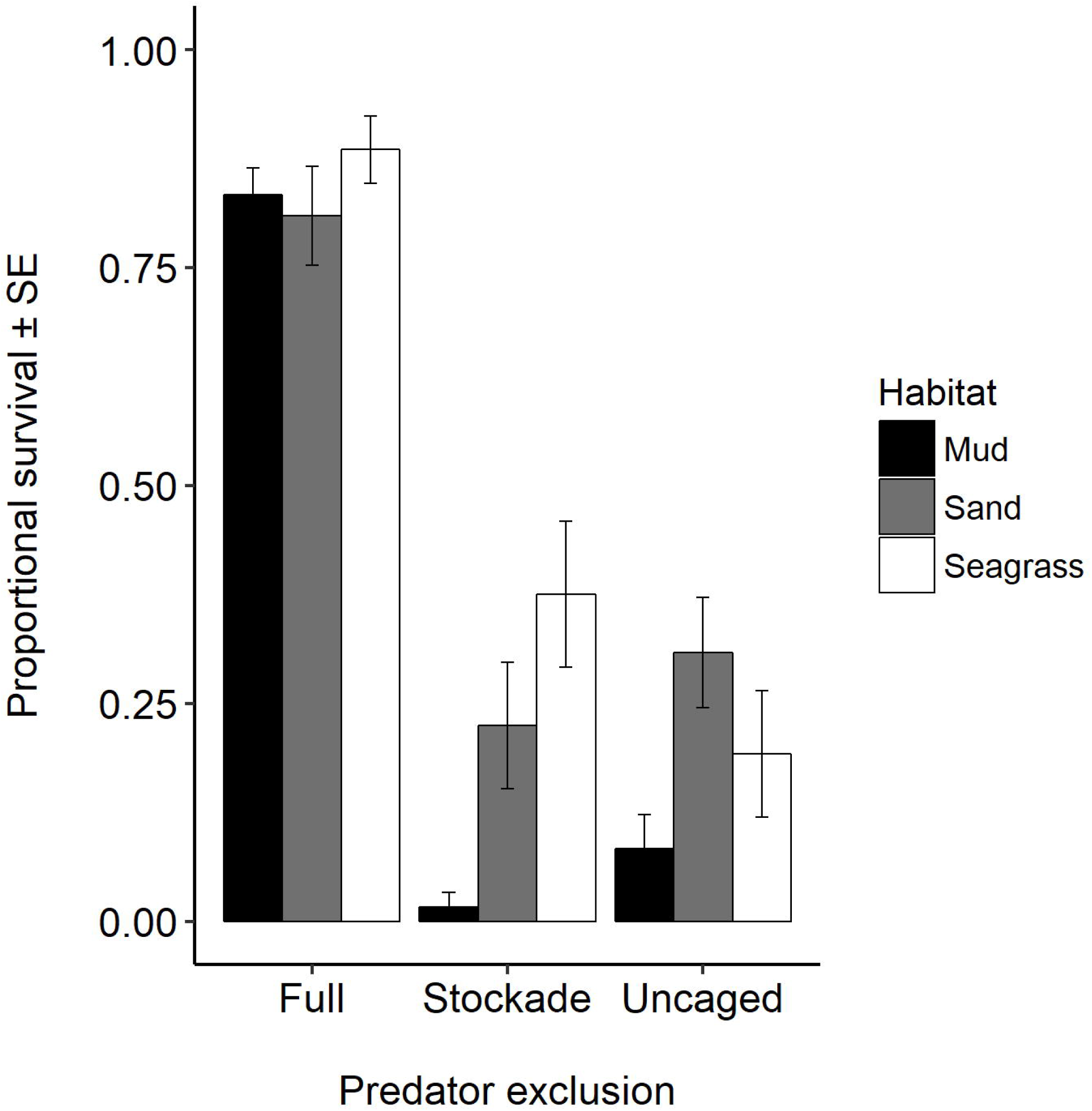
Behavior of blue crab *Callinectes sapidus* feeding on juvenile *Mya arenaria* and *Mercenaria mercenaria*. Shown are means (± 1 SE) of a) handling time (HT) for crabs feeding on *Mya*, b) HT for crabs feeding on *Mercenaria*, c) search time (ST) for crabs feeding on *Mya*, d) ST for crabs feeding on *Mercenaria*, e) encounter rate (ER) for crabs feeding on *Mya*, and f) ER for crabs feeding on *Mercenaria*. Lines of different colors and patterns represent different habitat types (shell = shell hash; oyster = oyster shell), and means were calculated from n = 3 trials.

Search time was shorter in low-density trials than in medium-density trials (Fig 4c, d; Table 2), but the effect of one main effect depended on the conditions of the others. The two treatments with the longest mean search times were *Mya* at medium density in seagrass (5.67 h) and *Mya* at medium density in oyster shell (5.56 h). The overall mean search times for *Mercenaria* at low and medium densities were 1.22 h and 1.91 h, respectively. The overall mean search times for *Mya* at low and medium densities were 0.89 h and 4.16 h, respectively. *Mya* at medium densities had longer search times than the other species x density combinations, driving a significant species x density interaction (Supp. Table 5). However, relatively long search times for medium densities of *Mya* only occurred in certain habitats (sand, oyster shell, and seagrass), resulting in a three-way interaction (Supp. Table 6).

Encounter rate was significantly lower in low-density trials than in medium-density trials (Fig 4e, f; Table 2). The two treatments with the highest mean encounter rates were *Mya* at medium density in sand (4.08 ind. h^-1^) and *Mya* at medium density in seagrass (3.23 ind. h^-1^). The overall mean encounter rates for *Mercenaria* at low and medium densities were 0.79 ind. h^-1^ and 1.80 ind. h^-1^, respectively. The overall mean encounter rates for *Mya* at low and medium densities were 0.81 ind. h^-1^ and 2.85 ind. h^-1^, respectively.

## DISCUSSION

Blue crabs were the main predators of *Mya* in all habitats we examined, with no significant difference between stockades and uncaged plots and high incidence of crushed shells, which is evidence of crab predation rather than another source of mortality [3]. This was in line with our hypothesis that crab predation would be important. Despite evidence in the literature that schooling rays can result in mass mortality of bivalves [38], and evidence from gut content analysis that cownose rays consume *Mya* [28], we did not observe evidence that cownose rays increased predation in uncaged plots relative to stockade plots during the time frame of our field experiment (May). These results were contrary to our hypothesis and indicate that over the time and spatial scale of this study, rays were not a major source of mortality for *Mya*.

Predation-related mortality was high for juvenile *Mya* that were not protected by a cage. Over a period of five days, exposure to predators decreased survival of juvenile *Mya* by 76.3% as compared to caged individuals. Clam survival was habitat dependent, and both sand and seagrass provided more refuge from predation than mud. *Mya arenaria* has previously been shown to achieve a low-density refuge in sand [14,21]; however, the results from the field caging experiment went against our hypothesis that the added complexity afforded by seagrass habitats provides an extended refuge for juvenile *Mya*. In the laboratory study, there was an effect of habitat on predator-related mortality only for *Mya*, which had lower survival in sand and seagrass than in shell hash or oyster shell habitats. However, in the case of a prey species that relies on achieving a low-density refuge for persistence, proportional survival may not be the best measure of success. Shell hash, oyster shell, and seagrass habitats had higher occurrence of trials with at least one clam remaining, which may be biologically meaningful. Habitat that allows survival of one or a few clams may maintain the low-density refuge for *Mya*.

Seagrass did not provide a refuge from predation for *Mya* in the field or in the laboratory experiment. However, seagrass in both studies was patchy; mesocosms were small, and caging sites were chosen so that the three habitat types (mud, sand, and seagrass) were in close proximity. Fragmented seagrass may not be able to provide much protection from generalist predators such as blue crabs, especially if they feed efficiently at patch edges [39]. Despite little evidence for patchy seagrass as a refuge from predation from this study, *Mya* are more likely to be found in seagrass than all other habitat types in the lower Chesapeake Bay [24]. This indicates that dense, contiguous seagrass stands may still provide a refuge from predation for *Mya*. Future research examining the effect of seagrass density or patch size on the survival of juvenile *Mya* is warranted.

Predators on *Mercenaria* (thick-shelled infaunal) and *Mya* (thin-shelled infaunal) had significantly different functional responses. Predators on *Mya* had a type III sigmoidal functional response, with a negative relationship between density and proportional survival, as has been seen in previous studies [14]. Predators on *Mercenaria* had a type II hyperbolic functional response, as has been seen previously [16], exhibiting either a positive relationship between density and proportional mortality or no density dependence, depending on the habitat. This difference is relevant to population dynamics and persistence of these two bivalve species because a type II functional response is unstable and can lead to local extinction of prey if they are driven to low densities, but a type III functional response may lead to prey persistence at low density [7,40]. The type II functional response of predators feeding on *Mercenaria* means this bivalve species must remain at relatively high densities to achieve population stability. Conversely, the type III functional response of predators feeding on *Mya* allows the species to persist, even at very low density.

The differences in functional response of predators feeding on *Mya* and *Mercenaria* were likely due to differences in predator behavior. Predators had shorter search time and encounter rate when prey were in low densities as compared to high densities, in agreement with our hypotheses, as predators appeared to give up foraging. At low densities, encounter rate did not differ between the two bivalve species, indicating blue crabs had less trouble finding deep-burrowing clams than we hypothesized. There was no evidence that blue crabs spent less time foraging in complex habitats or when exposed to deep-burrowing prey; on the contrary, blue crabs spent more time searching for *Mya* at medium densities than they did searching for *Mercenaria* at medium densities, indicating crabs may have a preference for *Mya* as prey. This tendency of blue crabs to pass up *Mercenaria* as prey may explain why handling times for *Mercenaria* were not significantly greater than handling times for *Mya*; while some crabs spent the extra time opening up the thick-shelled clams (*Mercenaria*), many predators also gave up without investing much time into the encounter.

Declines in complex habitat will likely lead to declines in thin-shelled species such as *Mya.* Oyster shell and shell hash provided juvenile *Mya* some protection from predation in mesocosm trials; however, in Chesapeake Bay, hard-bottom substrate, such as shell, is relatively uncommon [41]. Loss of many bivalves in the Bay, including oysters [42,43] and large-bodied clams [24,44,45], will make hard-bottom shell-hash habitat even more rare in the future. Seagrass has also experienced declines in the Chesapeake Bay [46], resulting in a decrease of many potential sources of highly complex benthic habitat in the Bay and a subsequent decrease in refuge for thin-shelled clams. *Mya* may retain a low-density refuge from predation even with the loss of structurally complex habitats, though a loss of habitat-mediated refuge may eventually result in clam densities that are not sustainable.

Loss of complex habitat in the Chesapeake Bay may have little impact on thick-shelled, infaunal bivalves such as *Mercenaria, Rangia cuneata,* and ark clams (*Noetia ponderosa* and *Anadara* spp.). We did not see an effect of habitat on *Mercenaria* survival in the current study, yet in previous research, *Mercenaria* had higher survival in crushed oyster shell habitats than in sand or mud [33]. This inconsistency is likely due to the use of larger clams in the current study (∼30 mm shell length) as compared to the previous study, which used clams 5-10 mm shell length [33]. Ontogenetic shifts in functional response may drive spatial distributions of hard-shelled bivalves in Chesapeake Bay, which are most dense in oyster shell habitats [47]. However, the effect of habitat on survival of recruits does not appear to impact population dynamics of large *Mercenaria*, which were present in multiple size classes throughout the year in lower Chesapeake Bay. Future research should examine whether complex habitat reduces blue crab encounter rates with small (< 10 mm) *Mercenaria* to determine the relationship between this species and complex habitat over its entire ontogeny.

### Relevance for conservation

Understanding the mechanism underlying bivalve refuges from predation is important in a changing world. Loss of structured habitat such as seagrass, mangroves, coral reefs, and oysters is occurring world-wide [48]. There is a current research need for models that can be used to forecast the impacts of global change, such as habitat loss, on predator-prey interactions [49]. We demonstrated that understanding the effect of habitat loss on predator-prey interactions is improved by understanding the mechanisms prey use to defend themselves against predators and the effects of prey density.

Nonlinear predator-prey dynamics can result in catastrophic changes and regime shifts [50,51]. An examination of the functional response is key in predicting the result of predator-prey interactions over time, and determining if a population crash can be expected in a food web, potentially leading to a regime shift. For instance, functional responses will be a major factor in determining whether a species driven to low abundance is likely to become locally extinct, or if it is likely to persist [19]. Documenting the functional response of bivalve species with a variety of different physical characteristics can help ecosystem managers decide on which species to focus conservation efforts, since species with a type II functional response are at higher risk of local extinction [52,53], and populations exhibiting a type III functional response are generally more stable over time [21,54,55].

A better understanding of density-dependent predator-prey interactions can be used to inform a variety of ecosystem management decisions. For example, functional responses can be used to determine a threshold density for reintroduction of endangered or depleted species [56], stock enhancement, [12,13], and pest control [57,58]. Effective bivalve seeding efforts that take into account predation may help restore marine bivalves, many of which have experienced severe declines in the recent past [42,43,59,60]. A better understanding of density-dependent predator-prey interactions will assist in the effort to maintain the integrity of marine trophic interactions and the viability of marine resources.

## ACKNOWLEDGMENTS

We gratefully acknowledge the assistance given by the students and staff of the Community Ecology and Marine Conservation Biology labs at the Virginia Institute of Marine Science. This paper is contribution number XXXX from the Virginia Institute of Marine Science, College of William & Mary.

## SUPPORTING INFORMATION

**S1 Table. Summary of Tukey HSD results for the caging study interaction term between habitat and cage type.** For each pairwise comparison, 95% confidence intervals (CI) and adjusted p values are presented. Data were Box-Cox transformed (λ = 0.51) prior to analysis and are not back-transformed. Only interactions with significant p values at α = 0.20 are shown.

**S2 Table. Summary of Tukey HSD results for the mesocosm study proportional mortality interaction term between species and density.** For each pairwise comparison, 95% confidence intervals (CI) and adjusted p values are presented. Data were Box-Cox transformed (λ = -0.14) prior to analysis and are not back-transformed. Only interactions with significant p values at α = 0.20 are shown.

**S3 Table. Summary of Tukey HSD results for the mesocosm study bivalve proportional mortality interaction term between species and habitat.** For each pairwise comparison, 95% confidence intervals (CI) and adjusted p values are presented. Data were Box-Cox transformed (λ = -0.14) prior to analysis and are not back-transformed. Only interactions with significant p values at α = 0.20 are shown.

**S4 Table. Summary of Tukey HSD results for the mesocosm study *Callinectes sapidus* handling time interaction term between species and habitat.** For each pairwise comparison, 95% confidence intervals (CI) and adjusted p values are presented. Data were fourth-root transformed prior to analysis and are not back-transformed. Only interactions with significant p values at α = 0.20 are shown.

**S5 Table. Summary of Tukey HSD results for the mesocosm study *Callinectes sapidus* search time interaction term between species and density.** For each pairwise comparison, 95% confidence intervals (CI) and adjusted p values are presented. Data were fourth-root transformed prior to analysis and are not back-transformed. Only interactions with significant p values at α = 0.20 are shown.

**S6 Table. Summary of Tukey HSD results for the mesocosm study *Callinectes sapidus* search time interaction term between species, density, and habitat.** For each pairwise comparison, 95% confidence intervals (CI) and adjusted p values are presented. Data were fourth-root transformed prior to analysis and are not back-transformed. Only interactions with significant p values at α = 0.20 are shown.

